# Thermal niche warming is more consistent than range shifts in marine species under climate change

**DOI:** 10.64898/2026.02.18.706571

**Authors:** Federico Maioli, P. Daniël van Denderen, Max Lindmark, Marcel Montanyès, Eric J. Ward, Sean C. Anderson, Martin Lindegren

**Affiliations:** National Institute of Aquatic Resources, Technical University of Denmark (DTU), Henrik Dams Allé 202, Kongens Lyngby, 2800, Denmark; Department of Aquatic Resources, Swedish University of Agricultural Sciences (SLU), Turistgatan 5, Lysekil, 45330, Sweden; Conservation Biology Division, NOAA Fisheries, Northwest Fisheries Science Center, 2725 Montlake Blvd E, Seattle, WA 98112, USA; Pacific Biological Station, Fisheries and Oceans Canada, 3190 Hammond Bay Rd, Nanaimo, BC V9T 6N7, Canada; School of Resource and Environmental Management, Simon Fraser University, Burnaby, BC V5A 1S6, Canada.

**Keywords:** Marine climate change, Species redistribution, Multidimensional range shifts, Ocean warming, Demersal fishes, Thermal niche

## Abstract

Marine species are widely expected to shift poleward or into deeper waters in response to rising ocean temperatures. However, our knowledge is primarily based on studies examining range shifts along single dimensions at a time (e.g., latitude or depth). Failing to address how movements along multiple dimensions interact, including associated changes in thermal exposure, may result in misleading conclusions and predictions of species distribution and community composition under global warming. To address this knowledge gap, we here develop and apply a multidimensional framework that jointly evaluates climate-driven redistribution of marine fish across latitude, longitude, depth and realized thermal niches, based on long-term scientific bottom-trawl surveys throughout the North Atlantic and Northeast Pacific. Our results show that net redistributions are generally small and highly region-specific, while the realized thermal niches of species have warmed substantially over the past three decades. These findings demonstrate that spatial redistribution is generally failing to keep pace with rising temperatures and challenge the prevailing assumption that marine species will move to escape warming. This has direct implications for biodiversity indicators that rely on distributional shifts as evidence of climate impacts, as well as climate-informed management and conservation of marine ecosystems, fisheries, and biodiversity at large.

## 1 Introduction

The world’s oceans are warming rapidly, driving a major reorganization of marine biodiversity with consequences for ecosystems and human use [1–4]. Marine species are widely expected to respond to ocean warming by tracking their thermal niches—the temperature ranges suitable for growth, survival and reproduction [5]. Such tracking often involves shifts in geographic distribution [6]. Consistent with this expectation, studies across marine taxa and regions document widespread poleward and deeper movements, making these range shifts among the most frequently reported biological responses to climate change [7–10].

Despite widespread reports of poleward and deeper shifts, some populations remain stationary even under sustained warming, while others shift in unexpected directions [11–13]. These departures likely reflect the complex spatial structure of ocean warming, including heterogeneous climate velocities and physical constraints imposed by coastlines, bathymetry, and circulation patterns [14, 15]. They may also arise from biological variability among taxa, including differences in thermal physiology, habitat affinity, recruitment capacity, and food-web limitations [16–19]. What remains unclear is whether, once aggregated across species and regions, these diverse responses give rise to coherent net redistribution signal—such as poleward or deeper shifts—or whether no consistent pattern emerges under ocean warming.

Determining whether species responses combine into coherent redistribution patterns requires evaluating movement across multiple dimensions simultaneously. However, most previous studies examined redistribution along a single gradient at a time—most commonly latitude, or, less often, depth [20]. While these approaches have revealed important climate-related patterns and responses, they provide limited insight into how movements along different dimensions interact, and what this implies for net redistribution across regions. In marine systems, species may shift horizontally, move vertically into cooler waters, remain in place while tolerating greater thermal exposure, or combine these strategies [21]. Hence, without integrating these responses—including associated changes in realized niches—we lack a complete picture of climate-driven redistribution under ocean warming [13, 22].

To address this gap, we develop and apply a multidimensional framework that evaluates climate-driven redistribution across three complementary dimensions: horizontal range shifts (changes in range centroids), vertical redistributions (realized depth niches), and changes in realized thermal niches. Unlike previous efforts that have modeled these attributes separately, our novel approach allows us to examine correlations across these redistribution pathways. This framework allows us to assess not only the direction and magnitude of spatial movements, but also whether such movements effectively track shifting thermal environments. If populations readily track changing isotherms, their realized thermal niches should remain stable through time. Hence, systematic warming of realized niches indicate incomplete tracking and increasing thermal exposure. We apply this framework to decades of standardized scientific bottom-trawl surveys encompassing more than 200 well-sampled demersal (bottom-living) fish populations across 11 regions of the North Atlantic and Northeast Pacific. These regions span broad latitudinal and climatic gradients, include some of the fastest-warming continental shelf seas [23, 24], and contain some of the most comprehensive long-term records of marine biodiversity change (Fig. 1; [25, 26]). Using spatiotemporal and Bayesian mixed-effects models, we jointly estimate long-term trends in species’ horizontal distributions, depth niches, and realized thermal niches. By examining these responses both within and across regions, we assess whether species-level responses combine into coherent redistribution patterns across species and scales and evaluate the extent to which such movements reduce thermal exposure under ocean warming.

**Fig. 1:**
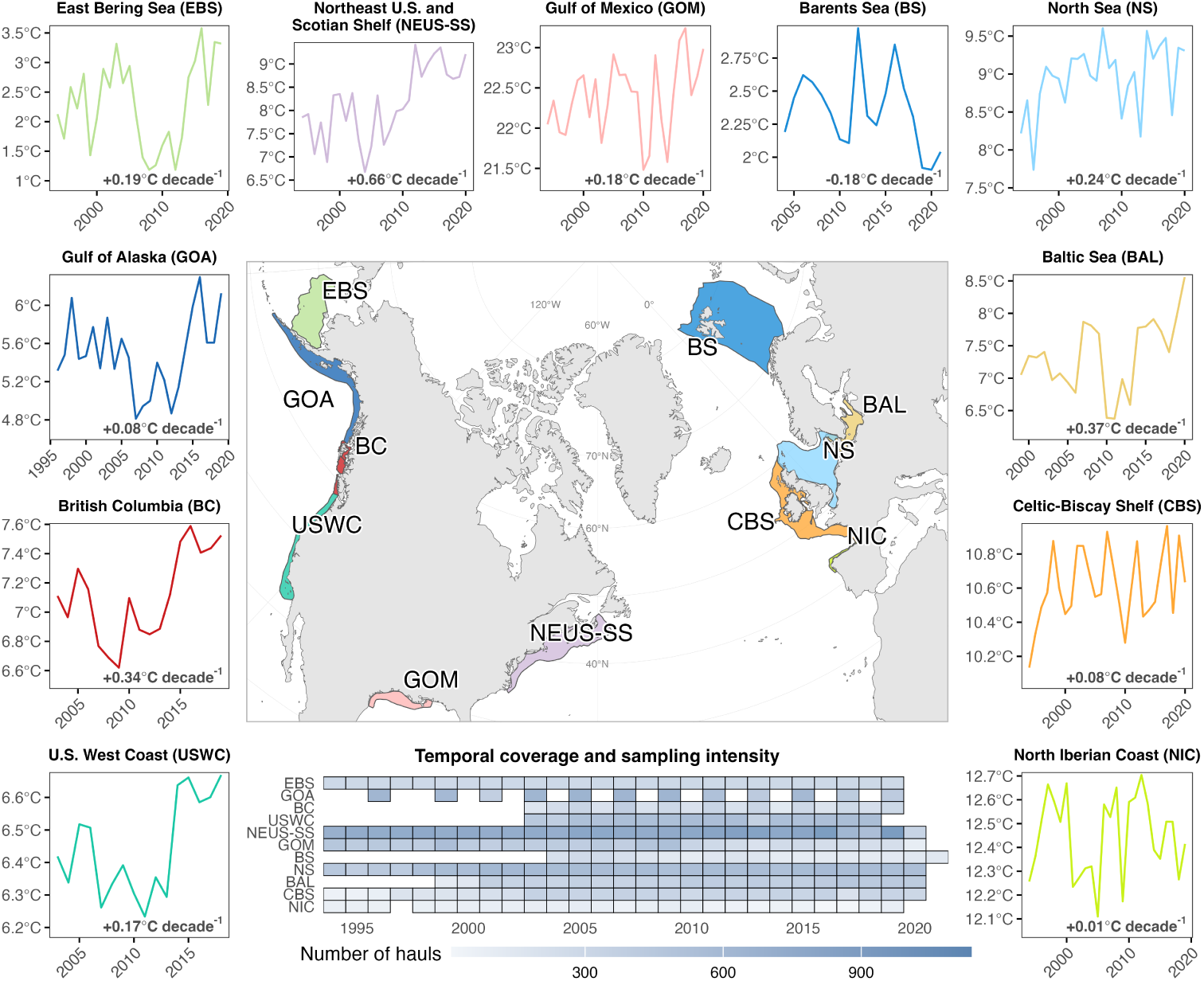
Overview of survey coverage and bottom temperature trends. The central map shows the regions included in the analysis, with colored polygons indicating region extents. The tile plot summarizes the number of hauls per region by year. Surrounding panels display regional trends in mean bottom temperature over time; bold values indicate the estimated decadal rate of temperature change.

## 2 Results

### 2.1 Limited net spatial shifts, but consistent niche warming

Overall, we detected no consistent directional shifts in either latitude or longitude across species and regions, as 95% credible intervals (CIs) greatly overlapped zero (Fig. 2a). This lack of net spatial redistribution reflects opposing species responses, with similar proportions of species shifting in opposite directions when aggregated across regions (Fig. 3a). The posterior median global slope (*β*_decade_) for latitudinal change was 1.33 km decade*^−^*^1^ [95% CI: -3.50, 7.59], and for longitudinal change -1.34 km decade*^−^*^1^ [95% CI: -6.70, 2.61]. Mean depth increased slightly over time, although uncertainty remained high (0.21 m decade*^−^*^1^ [95% CI: -0.49, 0.98]), while thermal niche temperatures increased consistently across species and regions (0.18 *^◦^*C decade*^−^*^1^ [95% CI: 0.06, 0.29]).

**Fig. 2:**
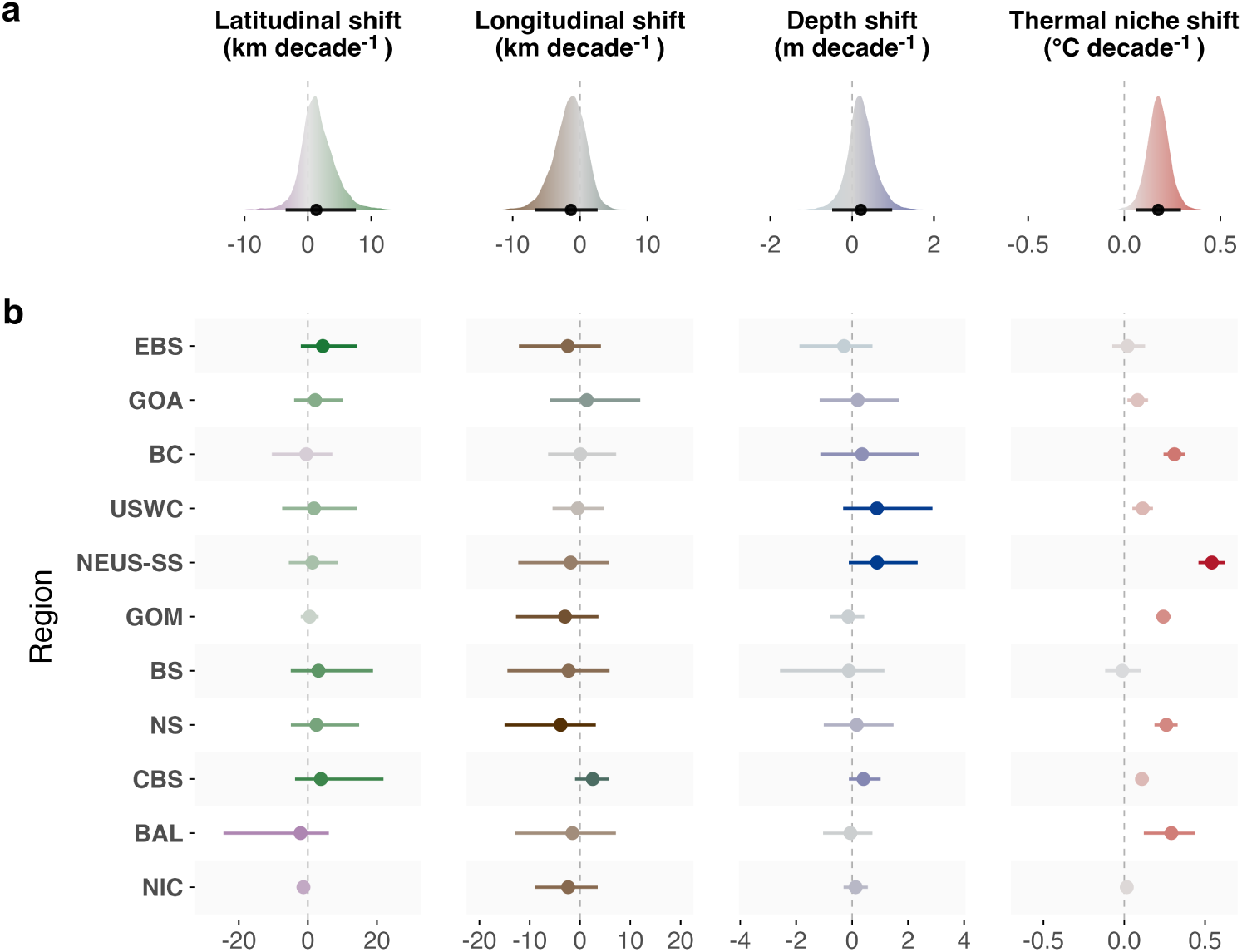
Trends in distributional shifts of demersal fish species, shown from left to right for latitudinal, longitudinal, depth, and thermal niche dimensions. Points and horizontal lines indicate posterior medians and 95% Bayesian credible intervals of decadal trends. (a) Global posterior slopes across all regions. (b) Region-specific posterior slopes. In (a), interval shading reflects the posterior distribution along a continuous gradient; in (b), color indicates median effect size, with more intense colors representing larger deviations from zero. Marine regions: EBS = Eastern Bering Sea, GOA = Gulf of Alaska, BC = British Columbia, USWC = U.S. West Coast, NEUS-SS = Northeast U.S. and Scotian Shelf, GOM = Gulf of Mexico, BS = Barents Sea, NS = North Sea, CBS = Celtic–Biscay Shelf, BAL = Baltic Sea, NIC = Northern Iberian Coast.

**Fig. 3:**
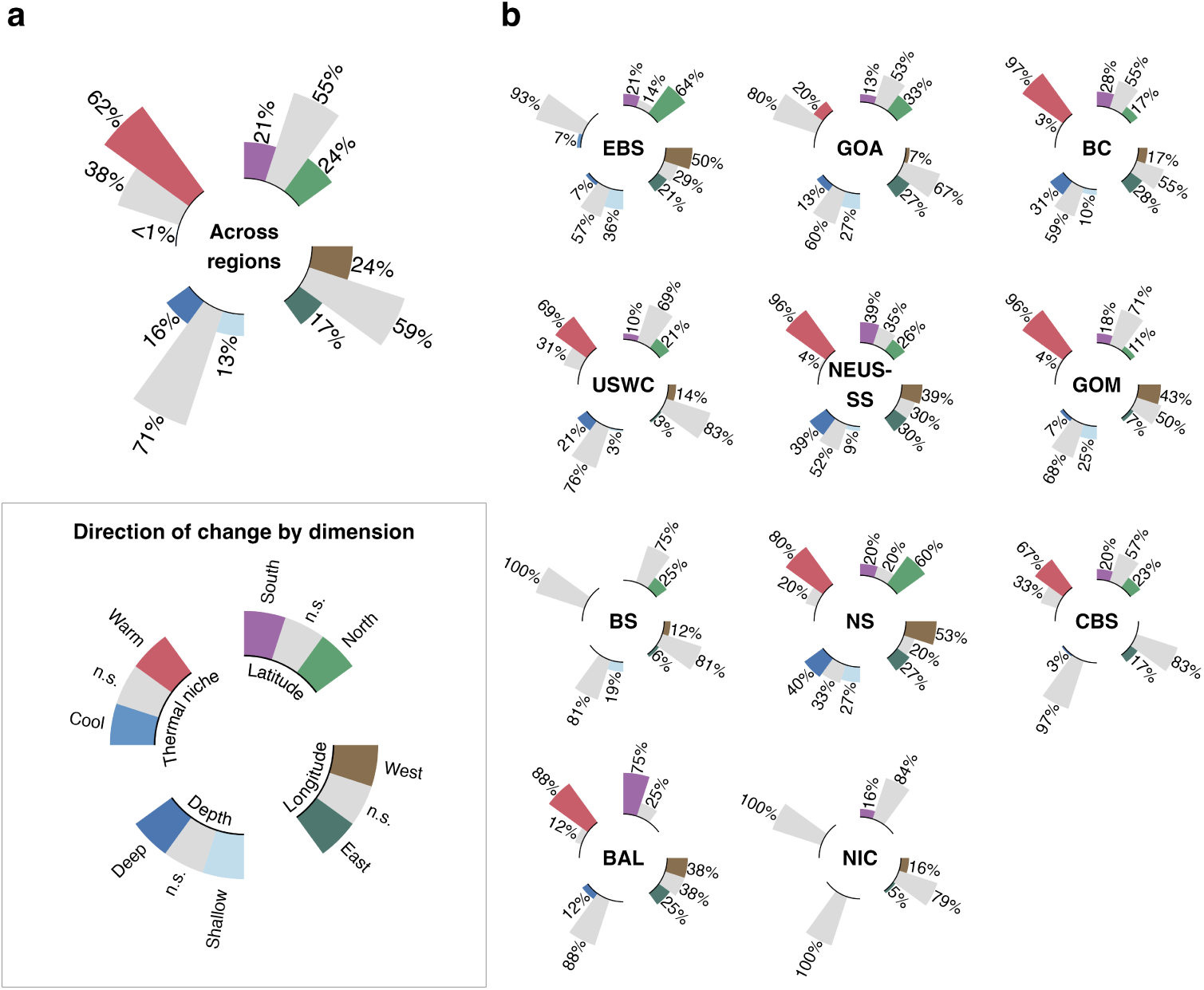
Proportion of species with supported shifts over time. (a) Across-region summary. (b) Regional patterns by marine region. Shifts are shown for latitude, longitude, depth, and thermal niche. Bars indicate the proportion of species with supported shifts (≥ 95% of the posterior distribution excluding zero); remaining responses are non-significant (n.s.). Marine regions: EBS = Eastern Bering Sea, GOA = Gulf of Alaska, BC = British Columbia, USWC = U.S. West Coast, NEUS-SS = Northeast U.S. and Scotian Shelf, GOM = Gulf of Mexico, BS = Barents Sea, NS = North Sea, CBS = Celtic–Biscay Shelf, BAL = Baltic Sea, NIC = Northern Iberian Coast.

### 2.2 Limited average spatial shifts, but consistent species responses within some regions

At the regional scale, posterior slopes for spatial shifts varied in direction and were generally small and uncertain, offering limited evidence for consistent changes in mean latitude, longitude, or depth within regions (Fig. 2b). Among spatial dimensions, depth showed the most consistent—though still modest—signals, with slight increases in mean depth along the U.S. West Coast (USWC; 0.88 m decade*^−^*^1^ [95% CI: -0.32, 2.87]) and the Northeast U.S. and Scotian Shelf (NEUS–SS; 0.89 m decade*^−^*^1^ [95% CI: -0.12, 2.34]). Although regional slopes were weak, many species within some regions shifted in the same direction (Fig. 3b). In the Eastern Bering Sea (EBS), 64% of species moved northward, with a similar proportion in the North Sea (NS; 60%), whereas 75% of species in the Baltic Sea (BAL) shifted southward. Depth shifts were also directionally consistent within some regions, with many species deepening in the Northeast U.S. and Scotian Shelf (39%) and British Columbia (BC; 31%), while relatively few shifted shallower. Depth shifts were absent or rare in other regions, with no significant changes detected in the Northern Iberian Coast (NIC) and only 3% of species shifting depth in the Celtic–Biscay Shelf (CBS).

Thermal-niche temperatures increased in nearly all regions and closely tracked local ocean-warming rates (Fig. 1; *ρ* = 0.92, *n* = 11, *p <* 0.001). Widespread thermal-niche warming was absent only in the Barents Sea (BS), Northern Iberian Coast, and Eastern Bering Sea. To isolate the effect of redistribution on thermal exposure, we contrasted realized thermal-niche trends with warming expected under static species distributions. Across most regions, realized and static warming were similar, suggesting that redistribution had limited net influence on thermal exposure (Supplementary Fig. S1). Within some regions, however,species shifting poleward or deeper experienced smaller increases in thermal-niche temperature than species that remained stationary or shifted in the opposite direction (Supplementary Fig. S2). Negative correlations between latitudinal shifts and thermal niche warming were strongest in the Northeast U.S. and Scotian Shelf (*ρ* = -0.88 [95% CI: -0.99, -0.67]), Celtic–Biscay Shelf (-0.84[95% CI: -0.98, -0.51]), North Sea (-0.80 [95% CI: -0.98, -0.44]), and Eastern Bering Sea (-0.62 [95% CI: -0.94, 0.01]). Depth shifts showed similar patterns in the Gulf of Mexico (-0.79 [95% CI: -0.98, -0.38]), British Columbia (-0.70 [95% CI: -0.96, -0.11]), and North Sea (-0.68 [95% CI: -0.95, -0.21]). By contrast, longitudinal shifts were generally weakly associated to thermal-niche change, except in the Northeast U.S. and Scotian Shelf, where longitudinal movements co-varied with latitude (0.86 [95% CI: 0.68, 0.96]). Spatial shifts along different axes also co-varied in other regions. For example, latitudinal and longitudinal shifts were strongly positively associated in the Gulf of Mexico, and negatively associated in British Columbia, and U.S. West Coast, reflecting coast-parallel movements. In the North Sea, latitudinal and depth shifts were correlated, with northward movements coinciding with deepening (0.86 [95% CI: 0.61, 0.97]).

### 2.3 Species show heterogeneous spatial shifts but niche warming is widespread

At the population level, spatial shifts were highly variable in both direction and magnitude (Supplementary Table S1, Supplementary Fig. S3). About 45% of the 226 populations shifted in latitude, with northward and southward movements occurring in similar proportions (Fig. 3a). The most extreme northward shift was seen in *Molva molva* in Celtic–Biscay Shelf (196.9 km decade*^−^*^1^ [95% CI: 142.2, 243.0]), and the largest southward in *Atheresthes stomias* in U.S. West Coast (-106.7 km decade*^−^*^1^ [95% CI: -138.5, -76.8]). Longitudinal shifts occurred in 40% of species, with 24% moving westward and 17% eastward; the strongest eastward and westward shifts were *Harengula jaguana* in Gulf of Mexico (115.9 km decade*^−^*^1^ [95% CI: 89.7, 142.4]) and *Squalus acanthias* in Northeast U.S. and Scotian Shelf (-109.5 km decade*^−^*^1^ [95% CI: -139.6, -79.0]), respectively. Depth shifts were less frequent, with 16% of species moving deeper and 13% shallower. The strongest deepening occurred in *Squalus suckleyi* in British Columbia (26.6 m decade*^−^*^1^ [95% CI: 18.7, 34.9]), and the greatest shallowing in *Anoplopoma fimbria* in British Columbia (-23.1 m decade*^−^*^1^ [95% CI: -53.4, -3.0]). In contrast, most species experienced warming in their thermal niches (62%), led by *Squalus acanthias* in Northeast U.S. and Scotian Shelf (1.1 *^◦^*C decade*^−^*^1^ [95% CI: 0.9, 1.3]), and only three species showing signs of cooling.

## 3 Discussion

Marine biodiversity is reorganizing under ocean warming, and numerous studies have documented poleward and deeper shifts across taxa and regions [9, 10, 27, 28]. These patterns have often been interpreted as evidence for a pervasive, directional redistribution of marine life in response to climate change. Our results question this overall narrative as we found little evidence for consistent, net spatial redistribution along latitude, longitude, or depth when responses were analyzed jointly and aggregated across regions. Instead, species-level movements were found to be highly variable and often opposing across species and regions. This resulted in weak cross-regional signals of spatial redistribution, with coherence emerging only in some particular regions. At the same time, we observed widespread warming of species’ realized thermal niches across most taxa and regions.

This widespread warming of realized thermal niches represents the most consistent signal across regions and taxa. It indicates that most species experienced increasing temperatures in the habitats they occupied, even where spatial redistribution occurred. This finding suggests that redistribution rarely provided full thermal compensation under ocean warming. Rather than maintaining stable thermal conditions through uniform isotherm tracking, these patterns align with evidence that realized thermal niches are dynamic and adaptive under climate change [29, 30]. A comparison with warming expected under spatially fixed distributions further supports this interpretation: when averaged across species within regions, realized and static warming were similar in most systems, indicating limited net buffering of background ocean warming through redistribution. Only in a few regions, such as the Eastern Bering Sea, did mean realized warming fall well below static expectations, consistent with partial compensation at the regional scale. Nonetheless, thermal compensation did emerge within some regions where species that shifted poleward or deeper tended to experience smaller increases in thermal-niche temperature. This is evident in systems with strong latitudinal temperature gradients and few barriers to movement, such as the Northeast U.S. and Scotian Shelf, Celtic-Biscay Shelf, the North Sea and the East Bering Sea, where poleward movements were associated with reduced niche warming. In other regions, depth-mediated responses appeared more important. For example, in the Gulf of Mexico—where poleward redistribution is geographically constrained—and in British Columbia, shifts to greater depths were associated with reduced thermal-niche warming. The North Sea represents a particularly coherent case, with many species shifting in concert both poleward and deeper, reflecting the tightly coupled horizontal and vertical thermal gradients in the area [31]. These well-documented examples, such as North Sea cod, illustrate that effective thermal tracking can occur under specific geographic and environmental configurations (Supplementary Table S1; 28, 32), but such cases remain the exception rather than the rule in our analysis.

The lack of strong net spatial redistribution contrasts with several cross-regional syntheses that report substantial, directional range shifts. For example, Poloczanska et al. [9] estimated mean poleward movements of roughly 30–70 km decade*^−^*^1^ at species’ centroids and leading edges, with comparable rates reported in the BioShifts database [33]. By comparison, the cross-regional trends we estimate were one to two orders of magnitude smaller and lacked a consistent direction. However, weak net spatial redistribution does not imply a lack of biological response to ocean warming. Rather, it emerges from the diversity of redistribution pathways available to marine species and the constraints that shape them. Although temperature gradients broadly align with latitude and depth, local bathymetry, coastlines, ocean circulation, and habitat structure generate region-specific pathways or obstacles for movement [15, 34]. Furthermore, species traits, habitat associations, and non-thermal drivers such as fishing pressure, oxygen availability, and prey distributions may modulate these responses [16, 35–37]. As a result, heterogeneous—and sometimes opposing—species-level movements may arise within and among regions under similar warming trends, producing weak aggregate signals at broader spatial scales.

Our analysis focuses on well-sampled species from standardized surveys, enabling consistent estimation of spatial and thermal metrics and explicit propagation of uncertainty across multiple dimensions of redistribution and thermal niches. This design results in narrower taxonomic and geographic coverage than a global synthesis, but it allows redistribution pathways to be evaluated within a robust and unified statistical framework. Many cross-regional analyses pool heterogeneous datasets and rely on summary metrics such as range edges, often without explicitly accounting for uncertainty in underlying shifts [33, 38, 39]. Estimates of distributional change may also be sensitive to spatial variation in sampling intensity, survey coverage, and catchability if observation processes are not formally modelled [40]. By contrast, our two-step frame-work estimates spatial and thermal metrics directly from standardized survey data and models temporal trends jointly across multiple dimensions. This approach propagates uncertainty from observations to species-, region-, and cross-regional summaries and allows correlations among redistribution pathways to vary between regions. Recent work has shown that explicitly accounting for uncertainty can substantially shape conclusions about biodiversity status and trends, and our results extend this insight to spatial redistribution under climate change [41]. These methodological differences likely explain why we detect weaker net redistribution signals at cross-regional scales than earlier syntheses.

Taken together, our results show that marine species responses’ to climate change do not manifest as coherent redistribution to maintain a specific thermal niche. Instead, thermal niches tend to get warmer with climate change, and redistribution reflects multidimensional, context-dependent responses shaped by local environmental gradients, physical constraints, and species-specific ecological traits and habitat affinities. When aggregated across regions, these heterogeneous trajectories often offset one another, yielding weak net redistribution despite sustained ocean warming. This complexity challenges the prevailing assumption of uniform poleward and deepward tracking and calls for evaluating redistribution and thermal niche change jointly when assessing biodiversity responses to continued climate change.

## 4 Methods

### 4.1 Data sources

#### Fish observations

We compiled high-resolution data on fish biomass (kg km^-2^) from a large-scale collection of standardized, fishery-independent bottom-trawl surveys [26]. Each trawl deployment (haul) recorded the catch by species, with biomass standardized by the sampled (“swept”) area, alongside the haul’s location, depth, and time. We selected surveys with a consistent sampling protocol and at least 15 years of coverage, resulting in 17 surveys across 11 regions in the North Atlantic and Northeast Pacific (Fig. 1). To ensure robust estimates of species’ spatial and temporal dynamics, we kept only taxa that: (i) made up 99% of the cumulative regional biomass, (ii) occurred in at least 15% of hauls in that region, and (iii) were sampled in at least two hauls per year. These criteria yielded 226 fish populations (species–region combinations) across all regions (Supplementary Table S1).

#### Environmental data

We obtained bottom temperature data from the Copernicus Global Ocean Physics Reanalysis [42], which provides monthly estimates at 1/12*^◦^* (≈7 km) resolution. While in situ measurements exist for some surveys in certain years, we used this modeled product to obtain temporally averaged and spatially consistent time series. Bathymetric data were derived from the GEBCO 2023 dataset [43], which provides global seafloor depth at 0.00417° (≈400 m) resolution.

### 4.2 Spatiotemporal modeling

#### Model structure

We modeled species distributions and their temporal dynamics using spatiotemporal generalized linear mixed-effects models (GLMMs), a flexible class of hierarchical models widely used to quantify spatial and temporal variation in marine species distributions [40, 44]. For each species (Supplementary Table S1), biomass density was modeled with a Tweedie distribution, using year (as a factor) and log-transformed depth (modeled as a second-order polynomial) as predictors. To account for differences in sampling and seasonal variation we included survey identity and quarter as fixed effects where appropriate. Spatial and spatiotemporal variation was captured through random fields, approximated via the stochastic partial differential equation (SPDE) method with Gaussian Markov random fields [45]. Models were implemented in R with the sdmTMB package [46], which integrates finite-element meshes constructed with fmesher [47] into Template Model Builder (TMB) [48]. Full details on model structure, parameterization, and fitting procedures are provided in Appendix A.

#### Prediction grid and environmental matching

After model fitting and validation, we predicted species-specific biomass densities on a 4 × 4 km spatial grid (in local UTM coordinates) covering each region for all modelled years. For each grid cell, we assigned bathymetry and the average bottom temperature over the 12 months preceding the earliest survey month in a region, with temperature values extracted from the gridded product using nearest-neighbor assignment. This procedure generated a consistent spatiotemporal dataset of predicted biomass, depth, and temperature for downstream analyses.

#### Derivation of multidimensional spatial and thermal-niche metrics

To assess species responses to temperature change, we used the predicted biomass described above to compute annual metrics that characterize both spatial redistribution and thermal exposure of species, including: range centroids (UTM northing and easting), realized depth niche, and realized thermal niche. Horizontal shifts in distribution were quantified using range centroids rather than range edges, as centroids are less susceptible to noise and provide a clearer signal of overall distributional movement, thus offering a more stable and representative measure of geographic displacement [49]. Range centroids were calculated as the biomass-weighted mean UTM northing and easting (center of gravity) across grid cells [40]. The realized depth and thermal niches were computed as biomass-weighted averages of depth and bottom temperature, respectively, summarizing the environmental conditions primarily occupied by each species within each region (Table 1). To quantify the contribution of redistribution to thermal exposure, we additionally calculated a static thermal metric by holding biomass weights constant across years at their grid-cell mean. This metric represents the warming expected if species distributions had remained spatially fixed while ocean temperatures changed.

**Table 1:**
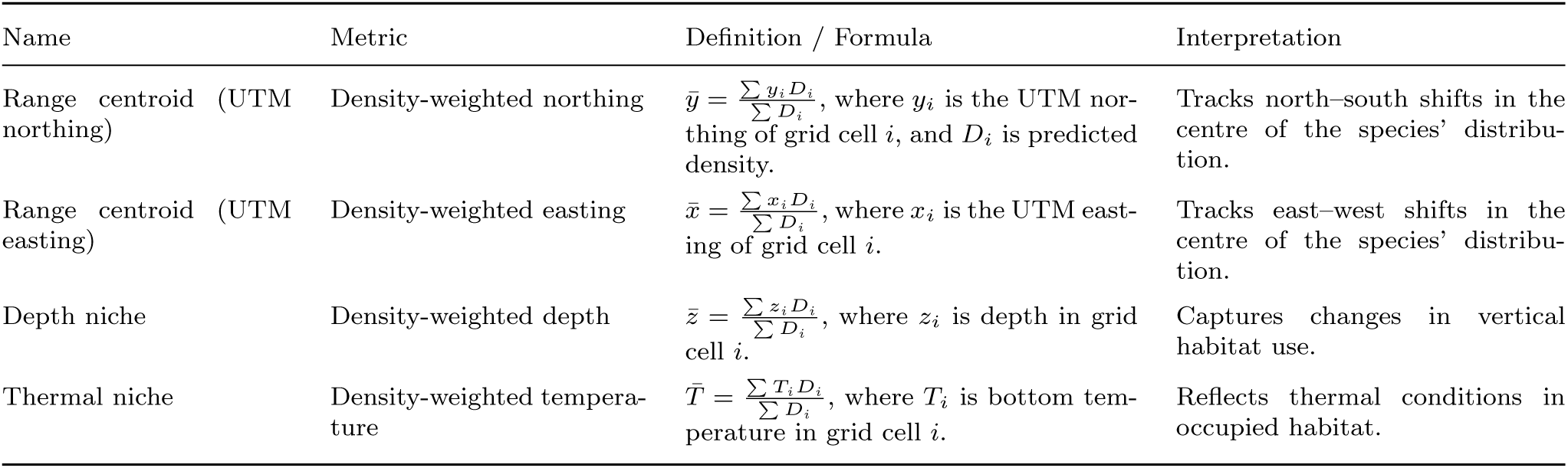
Summary of annual spatial and thermal metrics used to quantify species’ distributional and thermal responses to ocean warming.

### 4.3 Bayesian trend analysis

We estimated temporal trends in the derived spatial and thermal metrics using Bayesian mixed-effects models that jointly model multiple, potentially correlated responses. Each response was modeled using a Student-t likelihood to provide robust-ness to extreme values [e.g., 50], with uncertainty from the spatiotemporal modeling propagated via observation-specific standard errors. Correlations among responses were captured through a region-specific multivariate normal (MVN) covariance structure, allowing shared variation among spatial and thermal metrics to be estimated explicitly. Temporal trends were estimated hierarchically as varying slopes, with a global effect of time, region-level deviations, and species-specific deviations nested within regions, enabling inference across biological scales. To improve interpretability and facilitate model fitting, response variables and the time predictor were mean-centered within each species–region combination, such that their means equal zero. Under this parameterization, intercepts were omitted and all coefficients represent deviations from the species–region mean. Time was rescaled to decades, allowing slope parameters to be interpreted directly as rates of change per decade.

Thus the model can be written as:

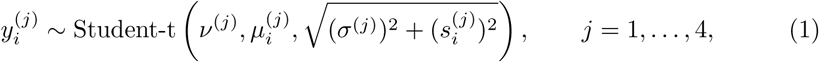

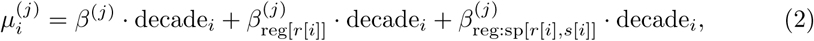

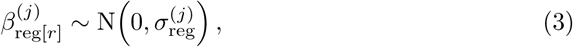

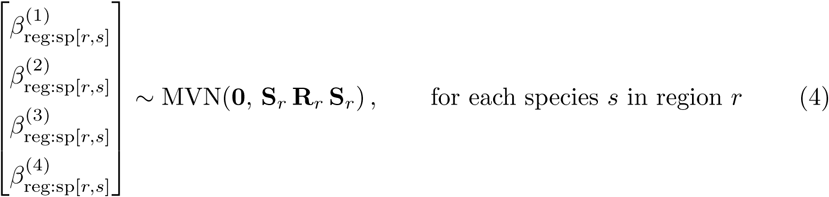

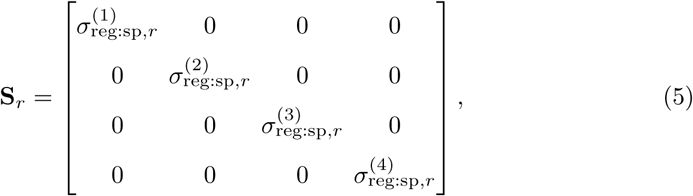

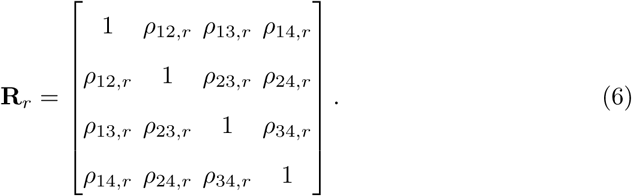

Here, *y_i_*^(*j*)^ denotes the response variable *j* for observation *i*, where *j* is indicates whether the variable is latitudinal centroid, longitudinal centroid, depth niche or thermal niche (Table 1). The term *s_i_*^(*j*)^ is the previously estimated standard error associated with observation *i*, *σ*^(*j*)^ is a response-specific residual scale parameter, and *ν*^(*j*)^ is the degrees of freedom. The linear predictor *µ_i_*^(*j*)^ (Eq. 2) models temporal trends as varying slopes, with a global effect of time *β*^(*j*)^, region-specific deviations *β*_reg*[r]*_^(*j*)^, and species-specific deviations nested within region *β*_reg:sp[*r*,*s*]_^(*j*)^.

Species-level slope deviation were modelled jointly across responses using a MVN distribution with region-specific covariance matrices **Σ*_r_*** = **S***_r_***R***_r_* **S***_r_*. This structure allows temporal trends in different response variables to be correlated among species within the same region, as controlled by **R***_r_*, while allowing these correlation patterns and the magnitude of response-specific variability to differ among regions.

Priors were weakly informative and guided by published rates of range shifts, depth changes, and ocean warming (Appendix B). We fit models in R using the brms package [51], which interfaces with Stan via rstan [52, 53]. Each model ran with four Markov chain Monte Carlo (MCMC) chains of 4,000 iterations, discarding the first 2,000 as warm-up. The remaining 2,000 samples per chain (8,000 total post–warm-up draws) formed the posterior distribution. Consistency with convergence was confirmed by R < 1.01, absence of divergent transitions, and effective sample sizes *>* 400 for all key parameters [54], as detailed in Appendix B.

## Supplementary information

Supplementary Table S1, Figs. 1–3 and Appendices A–B providing additional information on spatiotemporal models and Bayesian trend analysis.

## Acknowledgements

We thank all those who contributed to the collection of the survey data and the FISHGLOB consortium for providing public access to the dataset and for discussions held in Oldenburg. F.M. also thanks Christopher Griffiths for early discussions that helped shape this work.

## Data availability

All code used to perform the analyses and generate the figures is available at https://github.com/federico-maioli/species redistribution tracking and will be archived on Zenodo upon acceptance.

## Funding

F.M., P.V.D and M.L acknowledge support from the EU projects B-USEFUL (101059823) and ACTNOW (101060072). M.L. was supported by a research grant from the Swedish Research Council Formas (grant no. 2022-01511 to Max Lindmark).

**Fig. S1:**
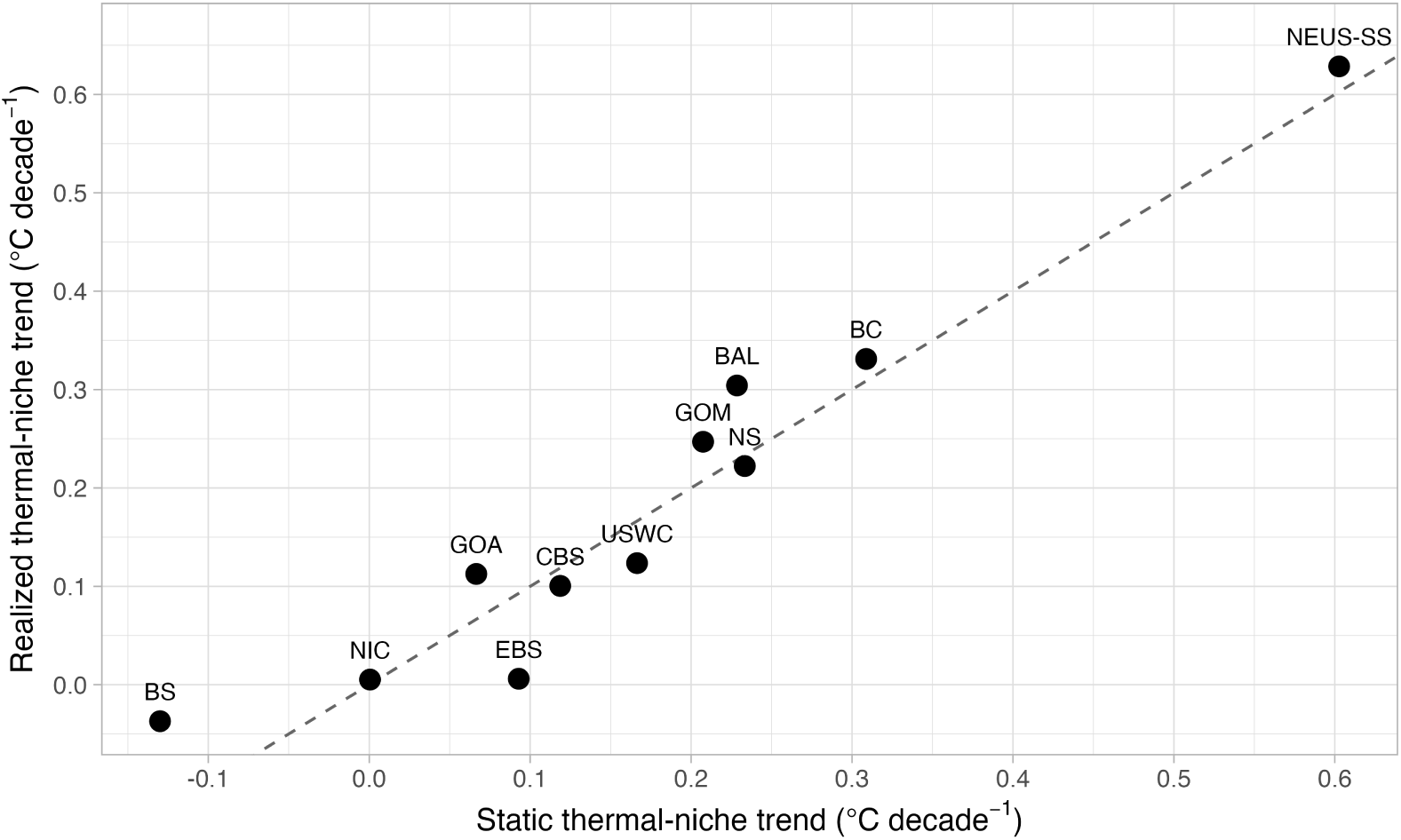
Realized versus static thermal-niche warming across regions. Mean decadal trends in realized thermal niches are plotted against warming expected under static species distributions (biomass weights held constant through time) for each region. The dashed 1:1 line indicates equal realized and static warming. Points below the line indicate regions where redistribution reduced thermal exposure relative to a fixed distribution (net buffering), whereas points above the line indicate greater realized warming than expected under static distributions. Marine regions: EBS = Eastern Bering Sea, GOA = Gulf of Alaska, BC = British Columbia, USWC = U.S. West Coast, NEUS–SS = Northeast U.S. and Scotian Shelf, GOM = Gulf of Mexico, BS = Barents Sea, NS = North Sea, CBS = Celtic–Biscay Shelf, BAL = Baltic Sea, NIC = Northern Iberian Coast.

**Fig. S2:**
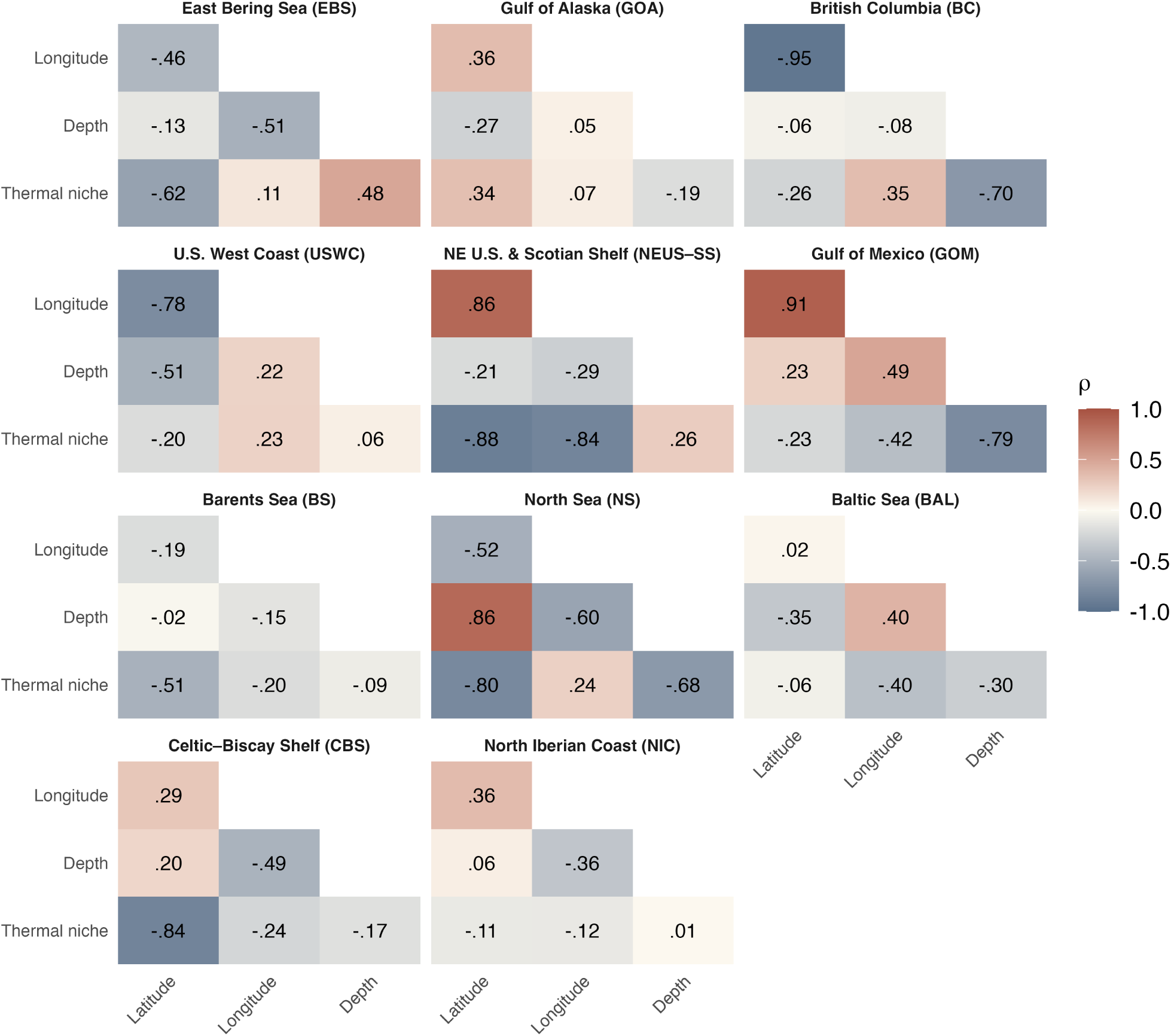
Heatmap of posterior mean correlations (*ρ_r_*) between trends in latitude, longitude, depth, and realized thermal niche across regions. Colors indicate the strength and direction of associations.

**Fig. S3:**
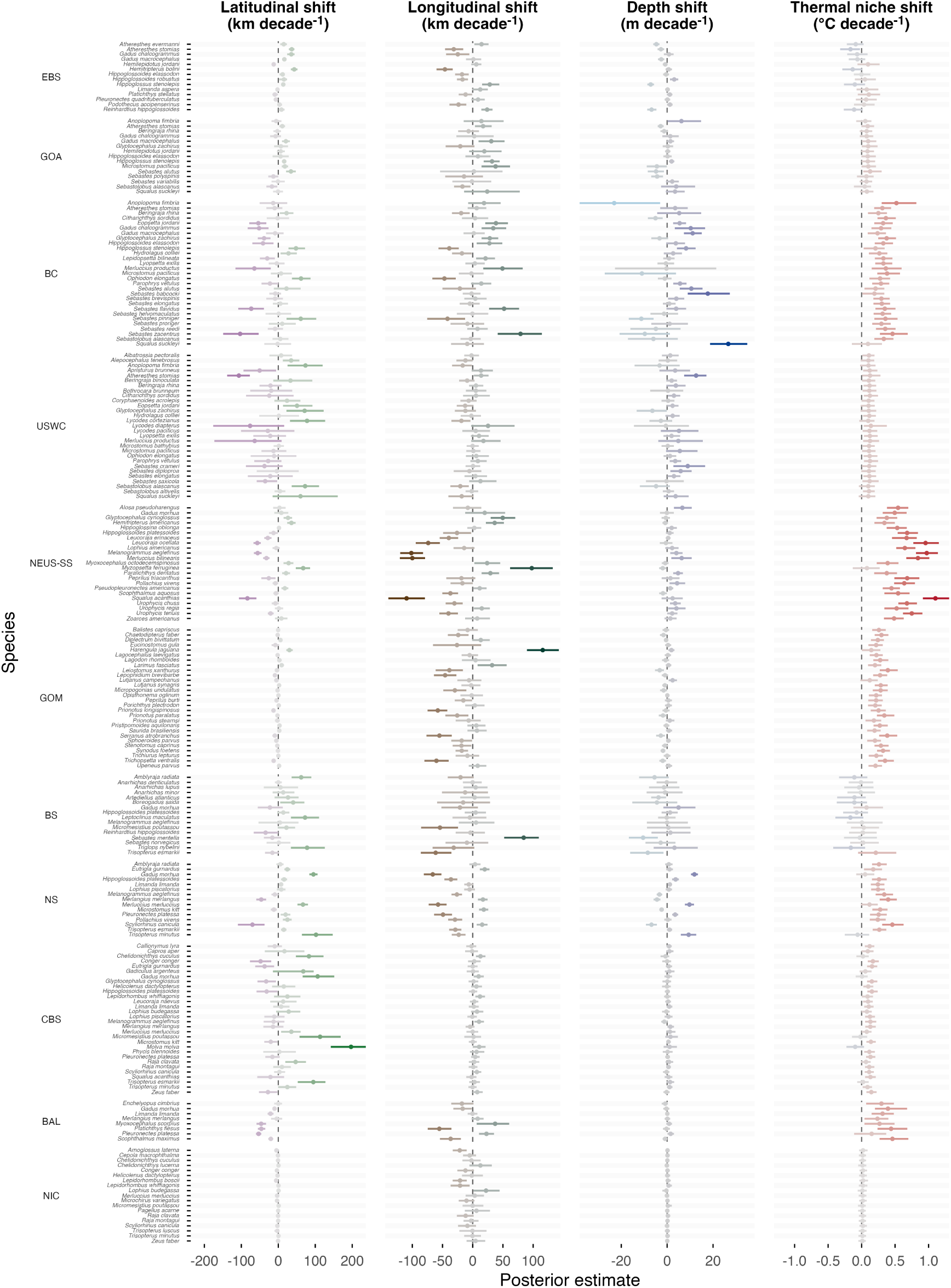
Species-specific trend shifts. Color shows median effect size; more intense colors indicate larger deviations from zero. Points and lines show median and 95% CI. Marine regions: EBS = Eastern Bering Sea, GOA = Gulf of Alaska, BC = British Columbia, USWC = U.S. West Coast, NEUS-SS = Northeast U.S. and Scotian Shelf, GOM = Gulf of Mexico, BS = Barents Sea, NS = North Sea, CBS = Celtic–Biscay Shelf, BAL = Baltic Sea, NIC = Northern Iberian Coast.

**Table S1:**
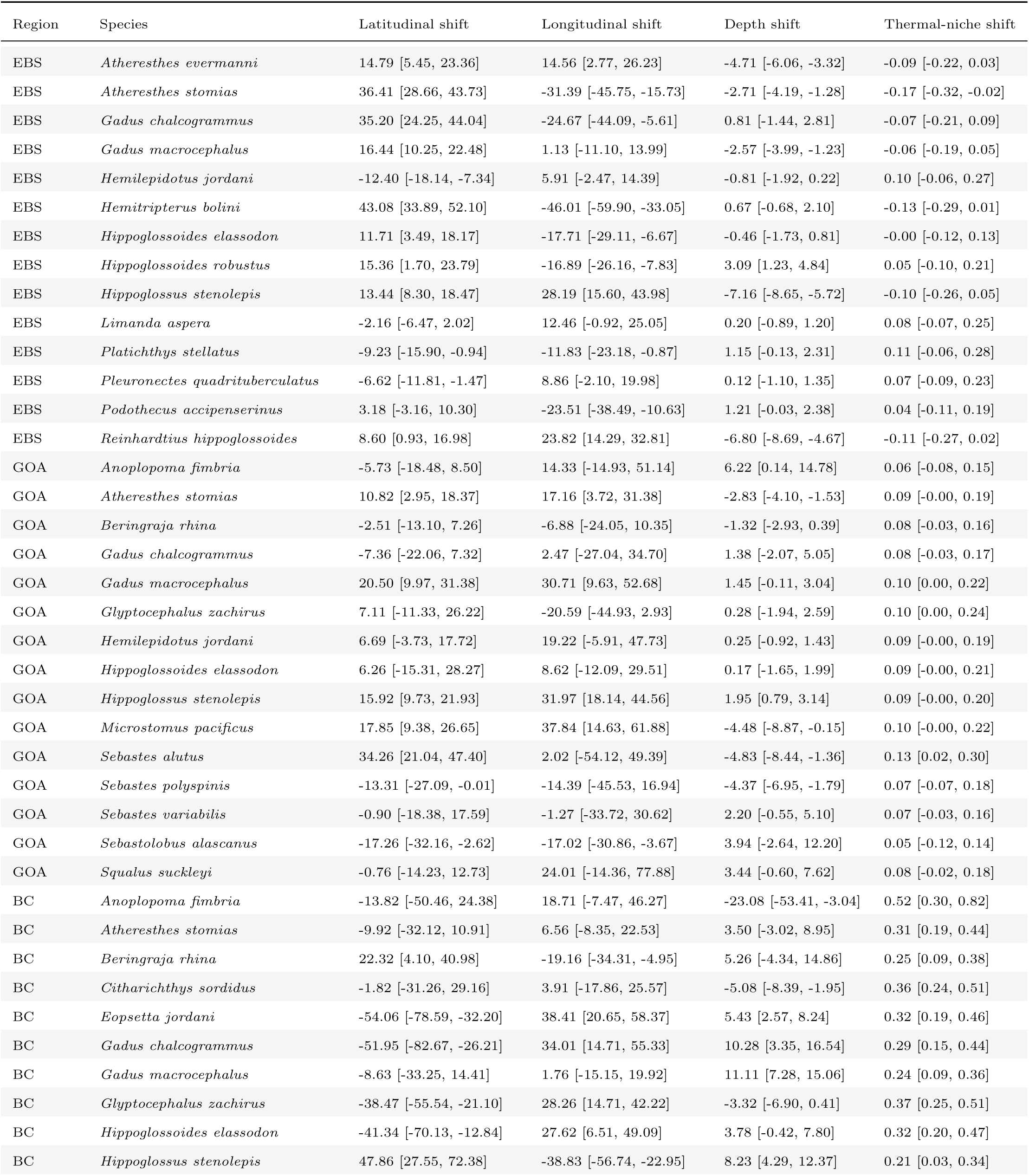
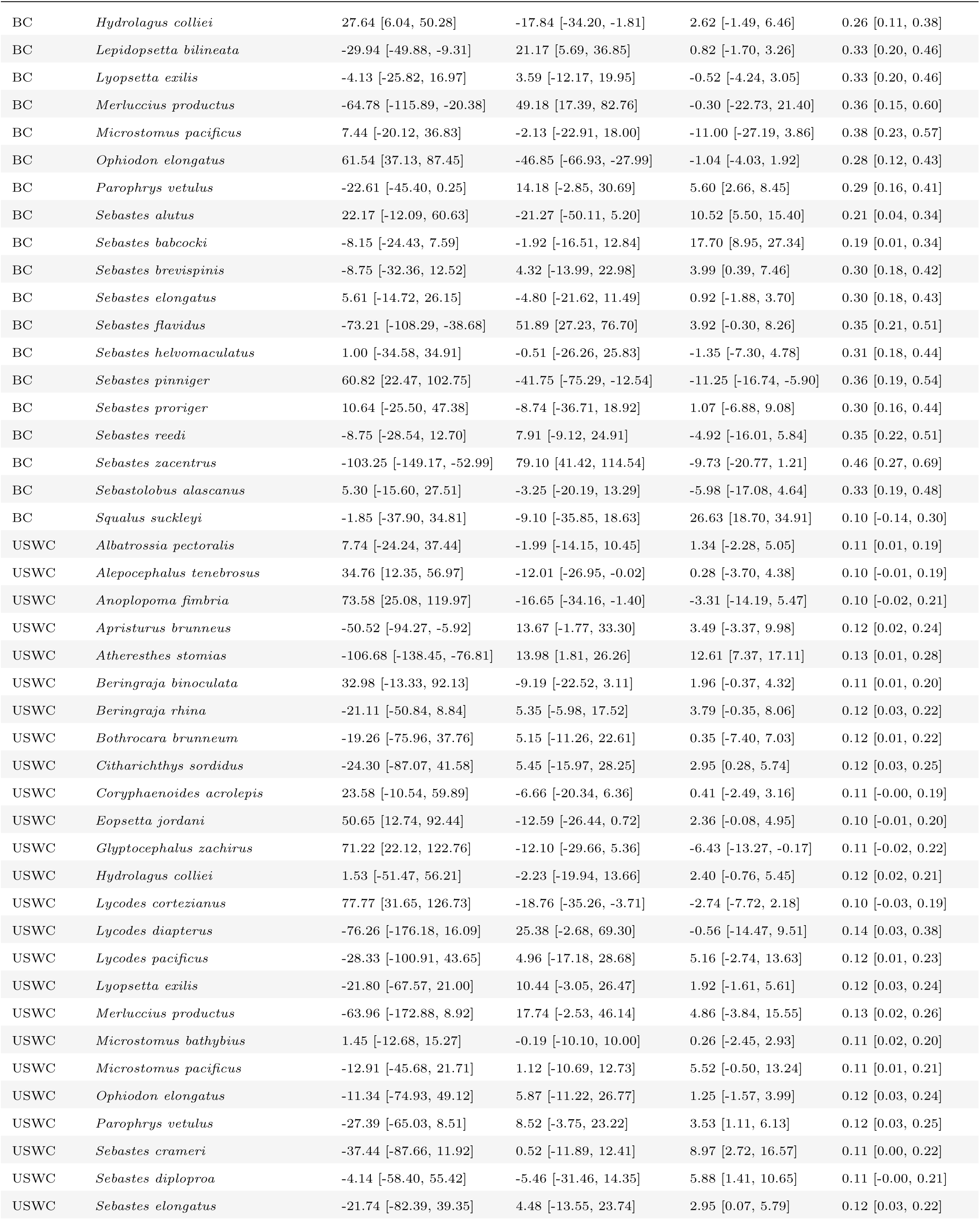
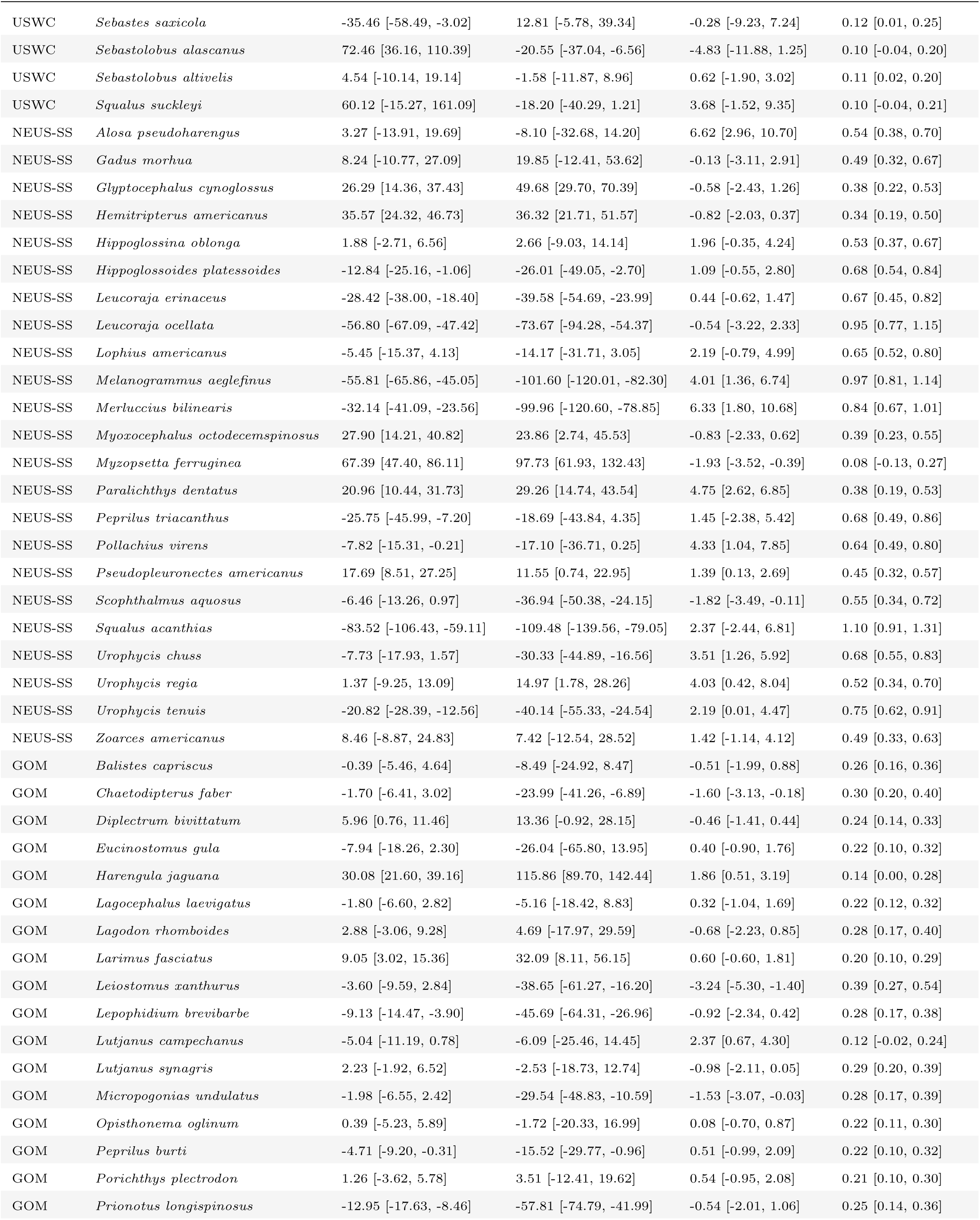
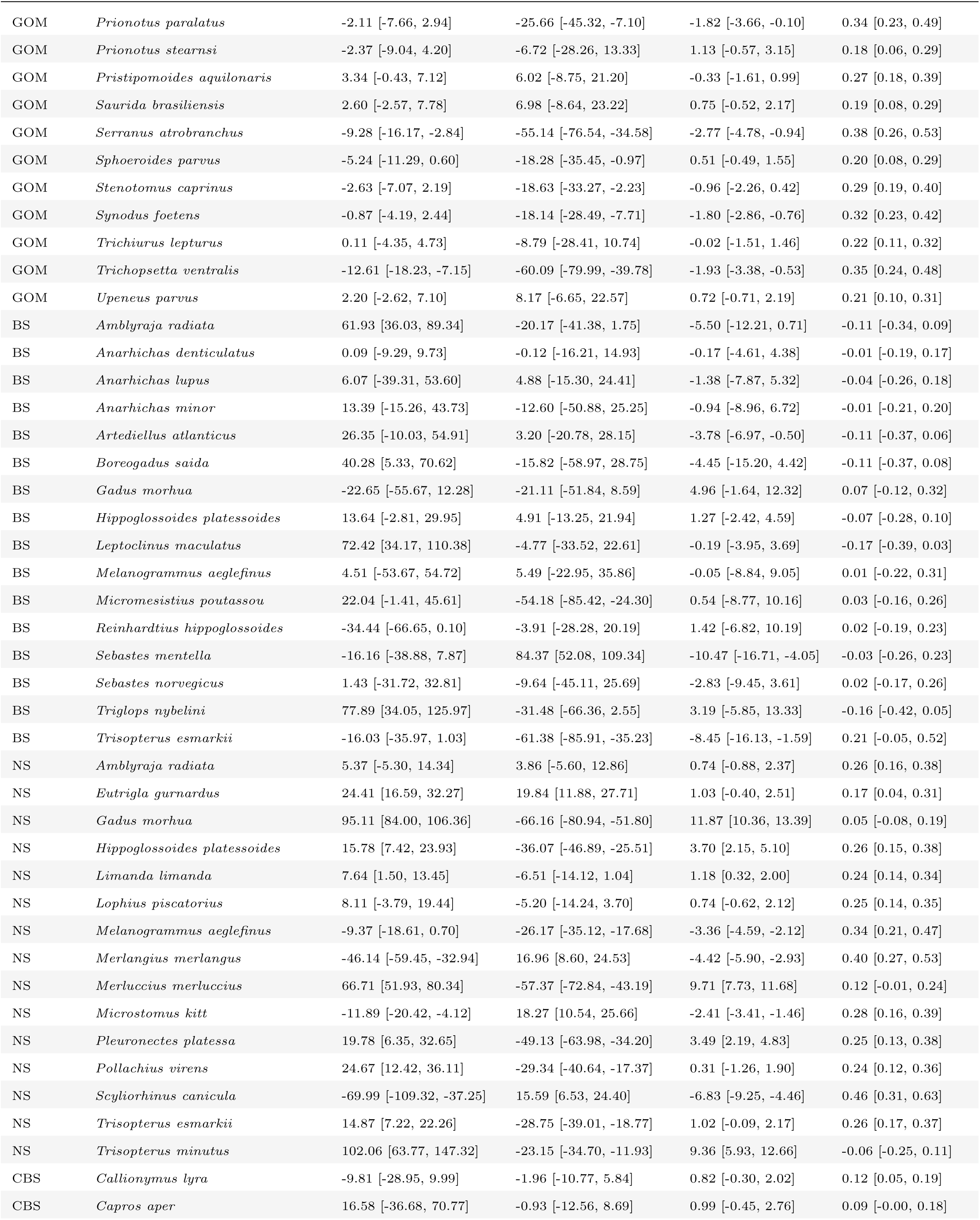
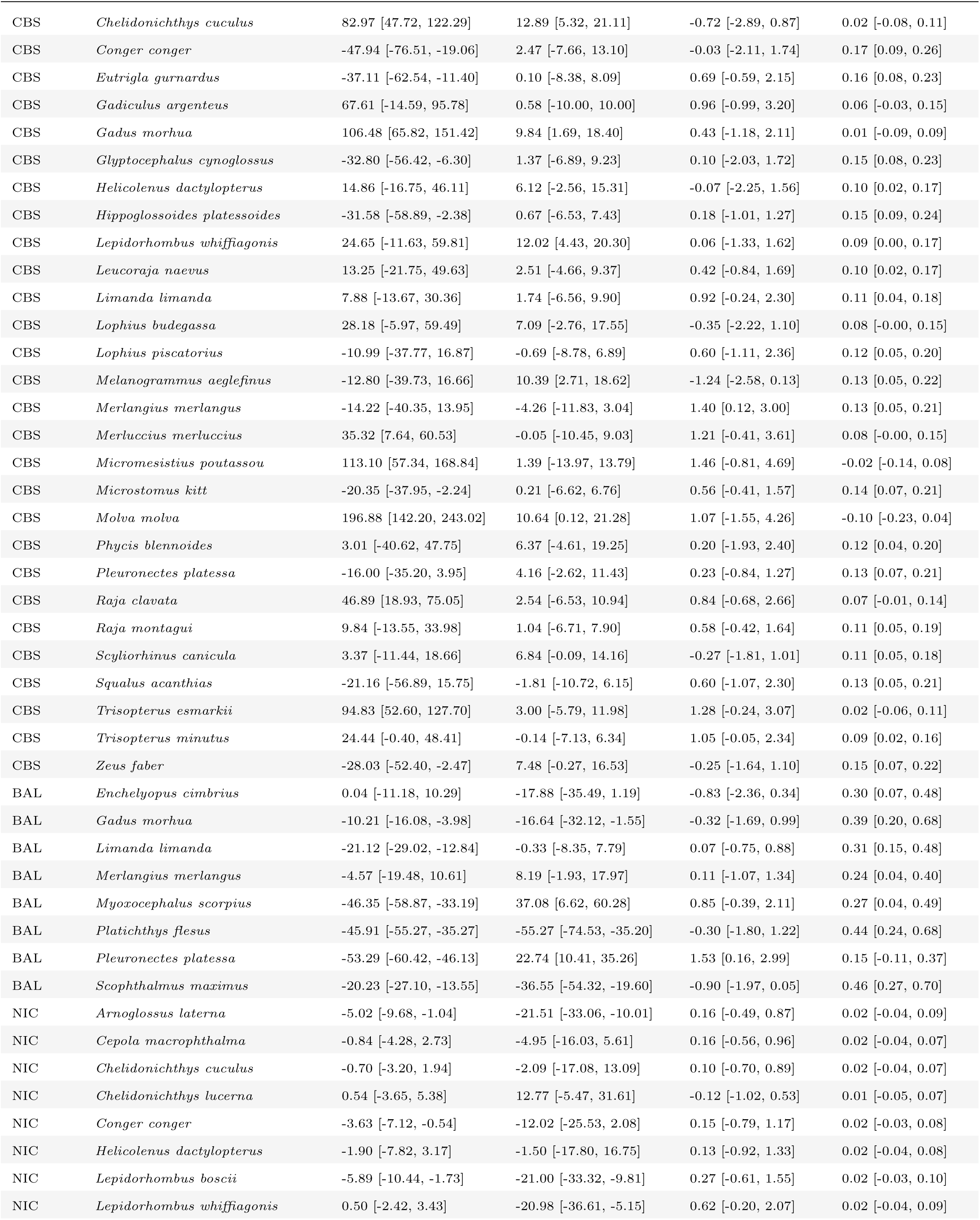
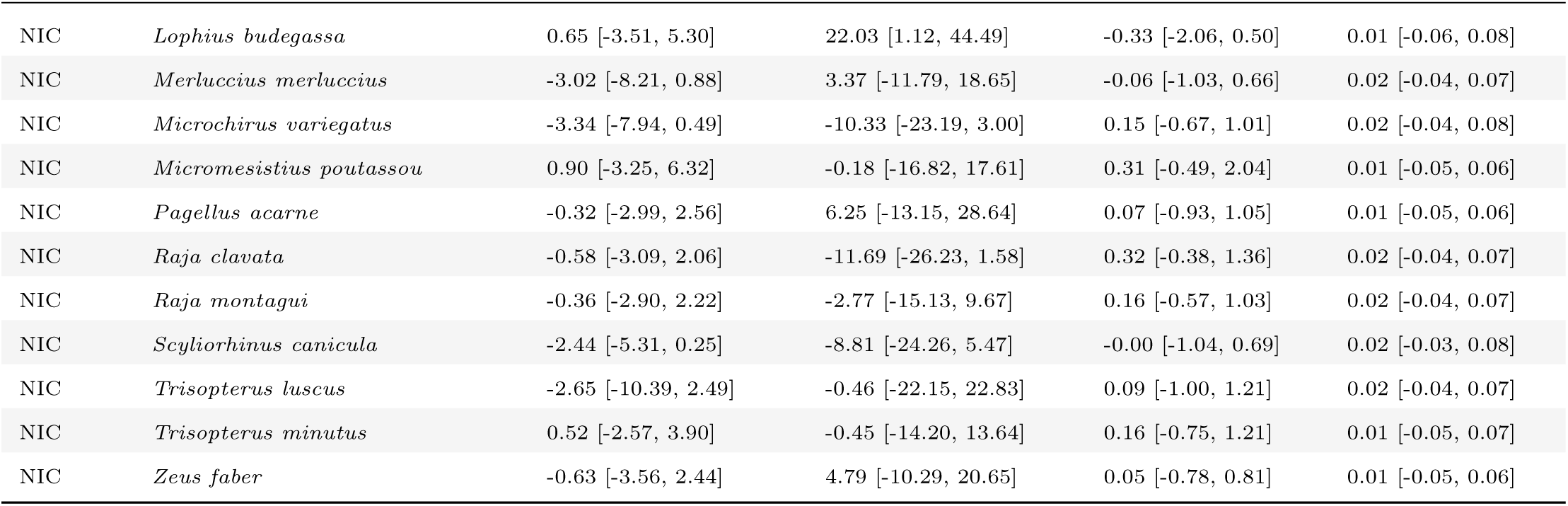
Posterior median decadal trends with 95% Bayesian credible intervals for latitudinal, longitudinal, depth, and realized thermal-niche shifts of demersal fish populations across regions. Latitudinal and longitudinal shifts are expressed in km decade*^−^*^1^, depth shifts in m decade*^−^*^1^, and thermal-niche shifts in *^◦^*C decade*^−^*^1^. Marine regions: EBS = Eastern Bering Sea, GOA = Gulf of Alaska, BC = British Columbia, USWC = U.S. West Coast, NEUS–SS = Northeast U.S. and Scotian Shelf, GOM = Gulf of Mexico, BS = Barents Sea, NS = North Sea, CBS = Celtic–Biscay Shelf, BAL = Baltic Sea, NIC = Northern Iberian Coast.

## Appendix A - Additional information on spatiotemporal models

The general form of the spatiotemporal GLMM can be represented as:

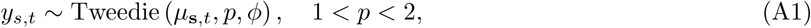

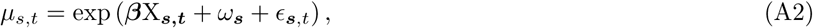

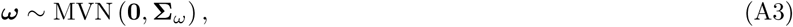

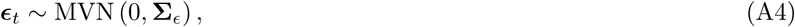

where *y***_s_***_,t_* represents fish density (kg km^-2^) at spatial location **s** and time *t*, *µ* is the expected mean density, and *p* and *ϕ* are the power and dispersion parameters of the Tweedie distribution, respectively. **X_s_***_,t_* is the design matrix of covariates including year (as a categorical variable) and a second-degree polynomial of log(depth), which helps constrain predictions to biologically plausible magnitudes across the depth gradient. The vector ***β*** contains the corresponding fixed-effect coefficients.

Because fish species are not constrained by survey boundaries, and some surveys cover only a small part of the distribution range of species we aggregated multiple surveys into broader regions for modeling when appropriate. Surveys with extensive and clearly defined coverage (e.g., the North Sea IBTS) were modeled independently, while those with contiguous spatial domains and partial overlap in time and space (i.e., surveys conducted in British Columbia, Northeast U.S & Scotian Shelf and Celtic-Biscay shelf) were combined (Fig A1). For regions with multiple surveys, we included survey identity as a fixed effect to control for differences in catchability. We did not apply this correction in British Columbia because surveys there follow harmonized field protocols and are typically analysed jointly [e.g., 1, 2]. In regions where surveys occur multiple times per year (BAL, NS, NEUS-SS, CBS, GOM), we included quarter as a fixed effect to account for within-year variation in sampling. In cases where a designated sampling period extended across multiple quarters, we used combined categories (e.g., Q2–Q3 or Q3-Q4).

The parameters *ω***_s_** and *ɛ***_s_***_,t_* (Equations A3–A4) represent spatial and spatiotemporal random effects, respectively, both modeled as Gaussian Markov random fields [3]. The spatial random field is shared across years, and represents static spatial effects such as habitat that are not accounted for by the fixed effects, while the spatiotemporal fields represent interannual spatial variation. We assumed spatiotemporal random fields to be independent across years to capture annual variation in spatial structure. While a first-order autoregressive (AR1) process could have been used to model temporal dependence, we opted for the independent formulation to strike a balance between computational efficiency and model flexibility. This approach allows spatial patterns to vary freely from year to year, accommodating potential non-stationary dynamics in species distributions without imposing strict temporal correlation structures. For British Columbia surveys, which have a biennial sampling structure, we modeled spatiotemporal variation as a random walk process and omitted the independent year fixed effects [2, 4]. This modifies Equation A4 into *ɛ****_s_****_,t_* ∼ MVN(*ɛ_t−_*_1_, **Σ***_ɛ_*) to allow for flexibility in estimating the spatial and temporal processes in years without data [2, 4].

Latent spatial and spatiotemporal random fields were approximated using a triangulated finite element mesh with a minimum spacing (cutoff) of 20 km, constructed with the fmesher R package [5]. This cutoff distance represents the smallest allowed distance between two mesh vertices. We assumed a shared range parameter (distance at which points are effectively independent [3]) between the spatial and spatiotemporal fields, while allowing each field to have its own variance.

Model parameters were estimated by maximizing the marginal log-likelihood using Template Model Builder (TMB; [6]), which applies the Laplace approximation to integrate out random effects. Models were fit in R 4.3.3 [7] using the sdmTMB package [8], which interfaces matrices from fmesher with TMB to implement the stochastic partial differential equation approach to approximating Gaussian random fields with Gaussian Markov random fields [3]. Only models with a positive-definite Hessian and a maximum absolute log-likelihood gradient *<* 0.001 with respect to all fixed effects were considered consistent with convergence and used for inference [8].

**Fig. A1:**
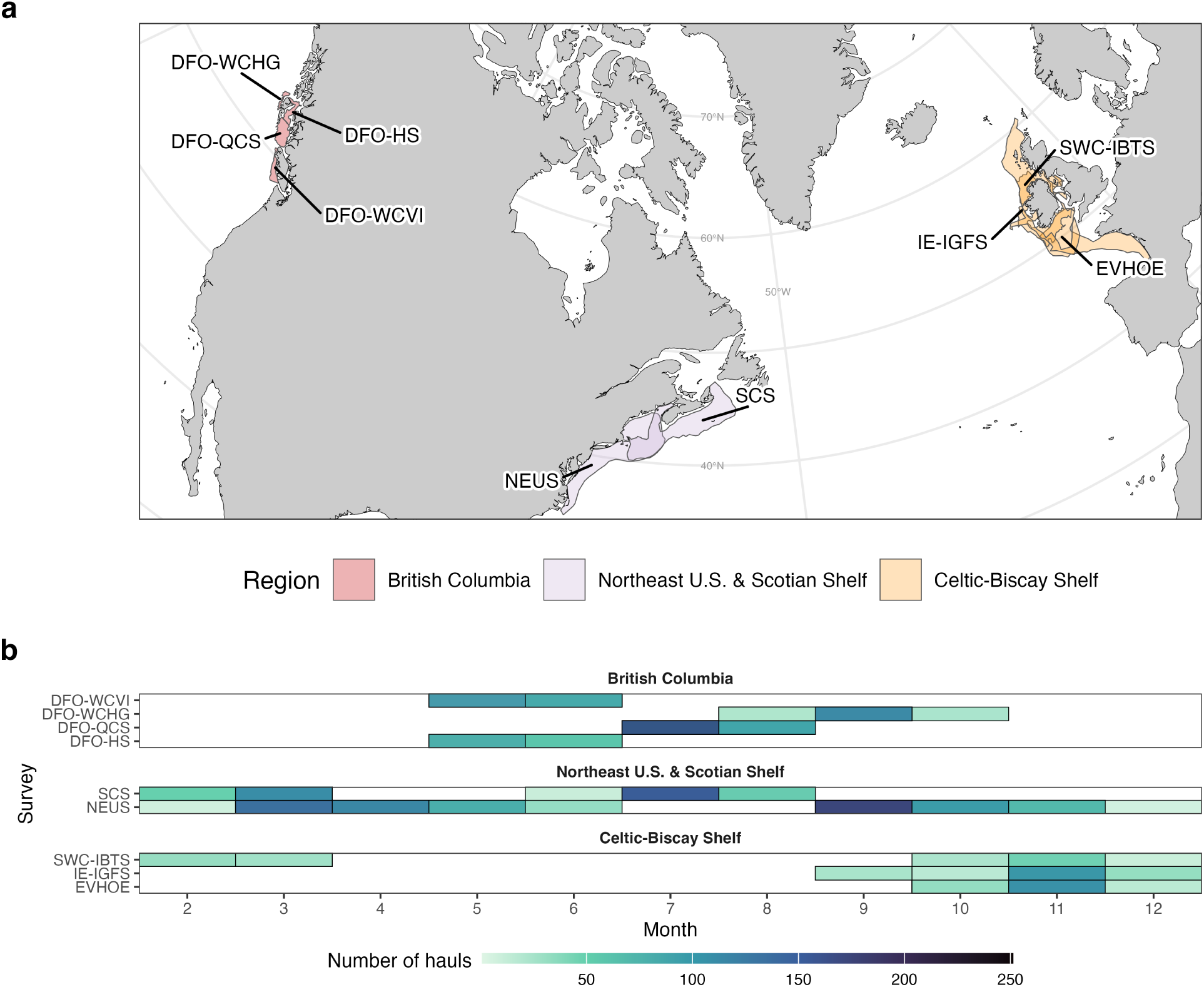
(a) Regions where multiple surveys were combined. (b) Overlap in survey months within each region.

## Appendix B - Additional information on Bayesian trend analysis

### Rationale for the Student-t likelihoods

We first modeled spatial and thermal responses with Gaussian likelihoods but posterior predictive checks indicated heavier tails than expected under normality. We therefore fit an alternative model using Student-t likelihoods, which better accommodates extreme values through an estimated degrees-of-freedom parameter. This specification improved predictive performance, as shown by higher expected log predictive density (ELPD) from leave-one-out cross-validation compared with the Gaussian model (Table B1; [1]). In both cases, correlations among responses were captured by modeling species-specific temporal slopes within regions as following a multivariate normal (MVN) distribution, allowing trends in different outcomes to vary jointly.

**Table B1:** Expected log predictive density (ELPD) differences between models. ELPD differences are reported relative to the best-performing model, which is assigned a value of zero. Negative values indicate poorer expected out-of-sample predictive performance. SE denotes the standard error of the ELPD difference

| Model | ELPD difference | SE difference |
| --- | --- | --- |
| Student-t + MVN | 0.00 | 0.00 |
| Gaussian + MVN | -2402.56 | 210.05 |

**Table B2:** Prior distributions for model parameters. Fixed effects (*β*), group-level and standard deviations (*σ*), degrees of freedom (*ν*) for the Student-t distribution, and the correlation matrix of region-level effects (**R***_r_*). The source column indicates whether the prior was changed from the default.

| Prior | Parameter | Coefficient | Group | Response | Source |
| --- | --- | --- | --- | --- | --- |
| LKJ(1) | $\mathbf{R}_r$ | | | | default |
| Normal(0, 50) | $\beta$ | decade | | lat centroid | user |
| Normal(0, 50) | $\beta$ | decade | | lon centroid | user |
| Normal(0, 5) | $\beta$ | decade | | depth niche | user |
| Normal(0, 0.5) | $\beta$ | decade | | thermal niche | user |
| Gamma(2, 0.1) | $\nu$ | | | lat centroid | default |
| Gamma(2, 0.1) | $\nu$ | | | lon centroid | default |
| Gamma(2, 0.1) | $\nu$ | | | depth niche | default |
| Gamma(2, 0.1) | $\nu$ | | | thermal niche | default |
| Student-t(3, 0, 31) | $\sigma$ | | | lat centroid | default |
| Student-t(3, 0, 27.9) | $\sigma$ | | | lon centroid | default |
| Student-t(3, 0, 3.9) | $\sigma$ | | | depth niche | default |
| Student-t(3, 0, 2.5) | $\sigma$ | | | thermal niche | default |
| Student-t(3, 0, 30) | $\sigma$ | decade | region | lat centroid | user |
| Student-t(3, 0, 30) | $\sigma$ | decade | region | lon centroid | user |
| Student-t(3, 0, 10) | $\sigma$ | decade | region | depth niche | user |
| Student-t(3, 0, 0.2) | $\sigma$ | decade | region | thermal niche | user |
| Student-t(3, 0, 40) | $\sigma$ | decade | region:species | lat centroid | user |
| Student-t(3, 0, 40) | $\sigma$ | decade | region:species | lon centroid | user |
| Student-t(3, 0, 20) | $\sigma$ | decade | region:species | depth niche | user |
| Student-t(3, 0, 0.4) | $\sigma$ | decade | region:species | thermal niche | user |
| Student-t(3, 0, 31) | $\sigma$ | | | lat centroid | default |
| Student-t(3, 0, 27.9) | $\sigma$ | | | lon centroid | default |
| Student-t(3, 0, 3.9) | $\sigma$ | | | depth niche | default |
| Student-t(3, 0, 2.5) | $\sigma$ | | | thermal niche | default |

### Hierarchical structure

An alternative to our approach is to fit separate models for each species and response. Such models, however, ignore information shared across taxa, regions, and outcomes. By using a hierarchical structure, we applied partial pooling: estimates borrow strength from the full dataset while still allowing region-and species-specific deviations. This shrinkage yields more stable estimates than independent fits (Fig. B1; [2]) and allows us to estimate global, regional, and species-level trends within a single unified framework. We modeled correlations among response variables at the regional level because species within the same region experience similar environmental conditions and constraints. This structure captures shared variation in temporal trends among species within regions while allowing species-specific departures from regional patterns.

### Priors specification and validation

All priors are summarized in Table B2. Throughout, Normal(*µ, σ*) and Student-t(*ν, µ, σ*) denote distributions with location *µ* and dispersion *σ*, where *σ* is the standard deviation for the Normal and the scale parameter for the Student-t, and *ν* denotes the degrees of freedom. We specified weakly informative priors for global temporal slopes (decade effects) based on published rates of range shifts, depth changes, and ocean warming (Fig. B2a). For latitudinal and longitudinal centroids, we used Normal(0, 50) km decade*^−^*^1^ priors, accommodating observed shifts of up to 30 km decade*^−^*^1^ in marine taxa [3]. For depth, we applied a Normal(0, 5) m decade*^−^*^1^ prior, consistent with reported deepening of 3.6 m decade*^−^*^1^ in North Sea demersal fish [4]. For thermal niches, we used a Normal(0, 0.5) *^◦^*C decade*^−^*^1^ prior, encompassing rates observed in rapidly warming shelf seas [5]. To model group-level variation in slopes, we assigned truncated Student-t distributions (0, ∞), with three degrees of freedom, allowing substantial but biologically plausible variation. Region-level slope standard deviations followed Student-t(3, 0, 30) km decade*^−^*^1^ priors for spatial centroids, Student-t(3, 0, 10) m decade*^−^*^1^ for depth, and Student-t(3, 0, 0.2) *^◦^*C decade*^−^*^1^ for thermal niches. For species nested within regions, we specified broader priors—Student-t(3, 0, 40) km decade*^−^*^1^, Student-t(3, 0, 20) m decade*^−^*^1^, and Student-t(3, 0, 0.4) *^◦^*C decade*^−^*^1^, respectively. These priors were consistent with the empirical variability observed in the data, providing appropriate regularization while remaining flexible enough to capture biologically plausible trends (Fig. B2b–c). All remaining model parameters retained the default priors provided by brms ([6]; Table B2).

### Model diagnostics

Model fit and convergence were assessed using posterior predictive checks (PPCs) and standard MCMC diagnostics. PPCs confirmed that the model captures the central tendency, variability, and distributional shape of the observed data (Fig. B3). Convergence diagnostics indicated *R*^^^ *<* 1.01 for all parameters, no divergent transitions, and effective sample sizes *>* 400, confirming reliable posterior sampling (Fig. B4; [7]). One parameter fell slightly below the recommended bulk ESS threshold, but visual inspection of the chains indicated satisfactory mixing (not shown), supporting reliable posterior sampling.

**Fig. B1:**
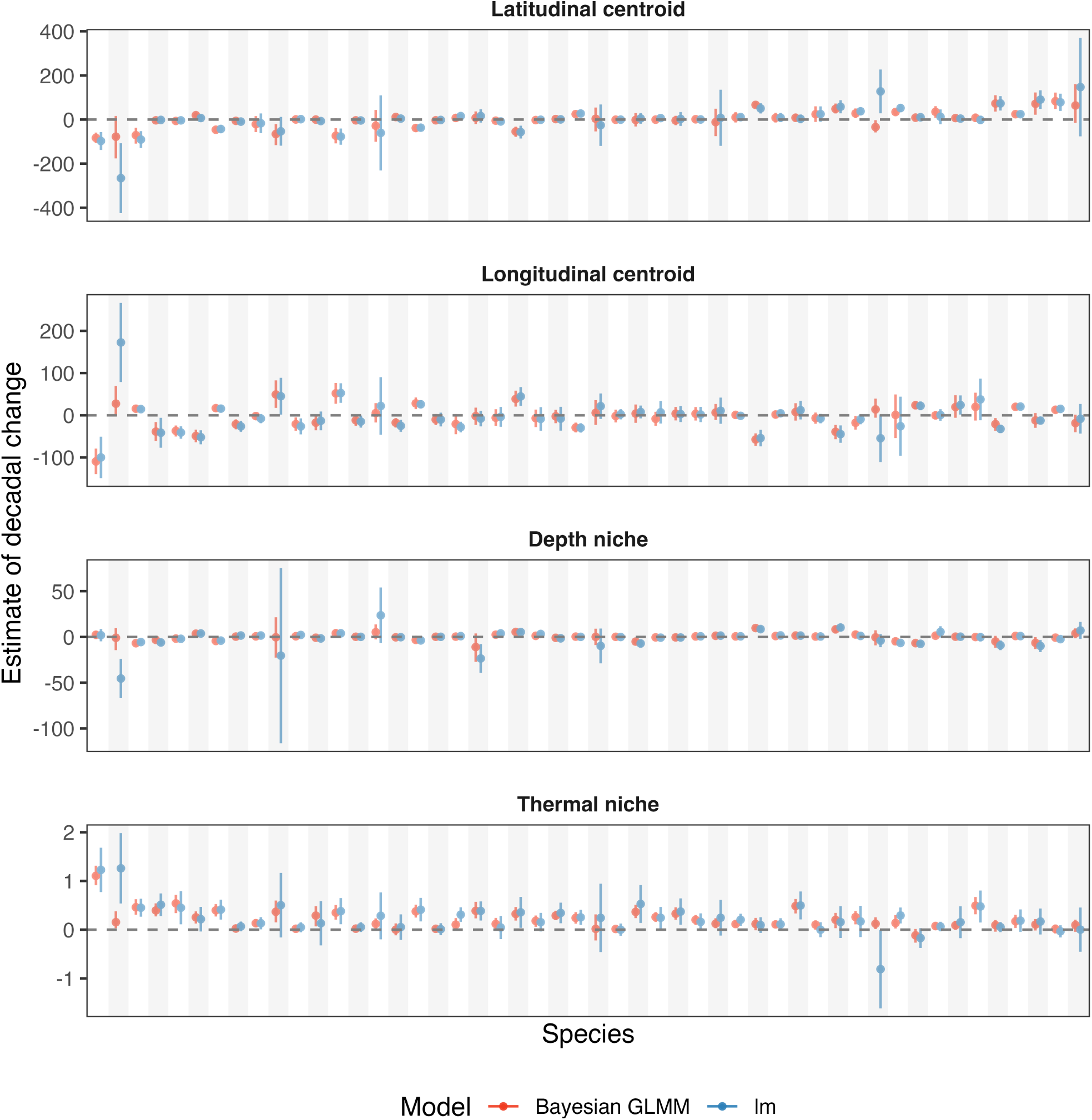
Comparison of hierarchical trend estimates versus separate linear trend fits. Posterior slopes from the full hierarchical model (red) are compared to slopes obtained from independent linear models (blue) fitted to a random subset of 50 species. Partial pooling in the hierarchical model shrinks extreme estimates toward the regional and global mean, producing more stable and biologically plausible trend estimates across species.

**Fig. B2:**
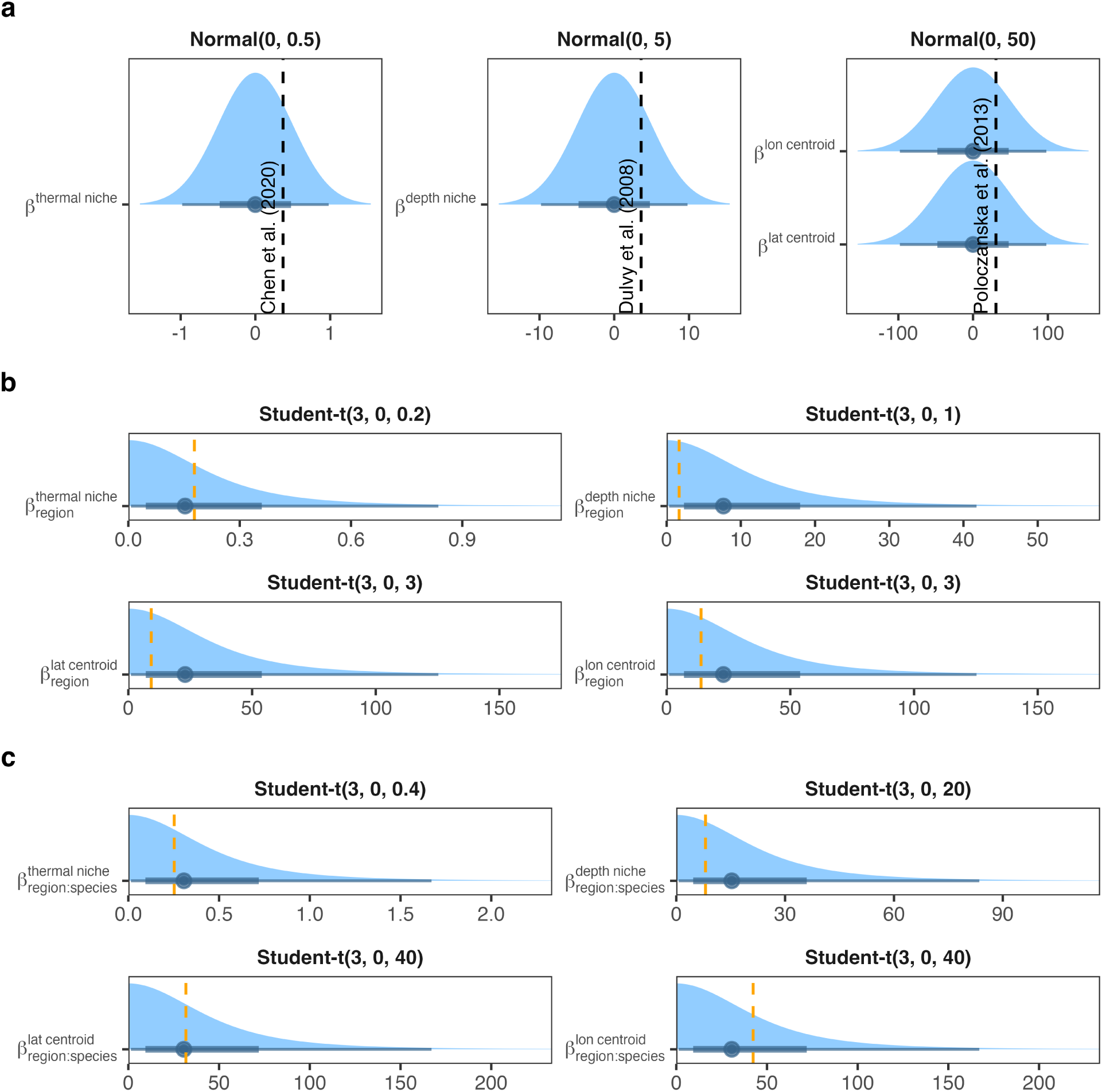
Prior specification and validation for temporal trend parameters. (a) Priors for global temporal slopes (decade effects) shown relative to literature-reported values (black dashed lines). (b) Priors for region-level slope variation compared with empirical distributions from the data (orange dashed lines). (c) Priors for species-level slope variation nested within regions compared with empirical distributions (orange dashed lines). Priors are weakly informative and consistent with observed variability, providing regularization without constraining plausible trends.

**Fig. B3:**
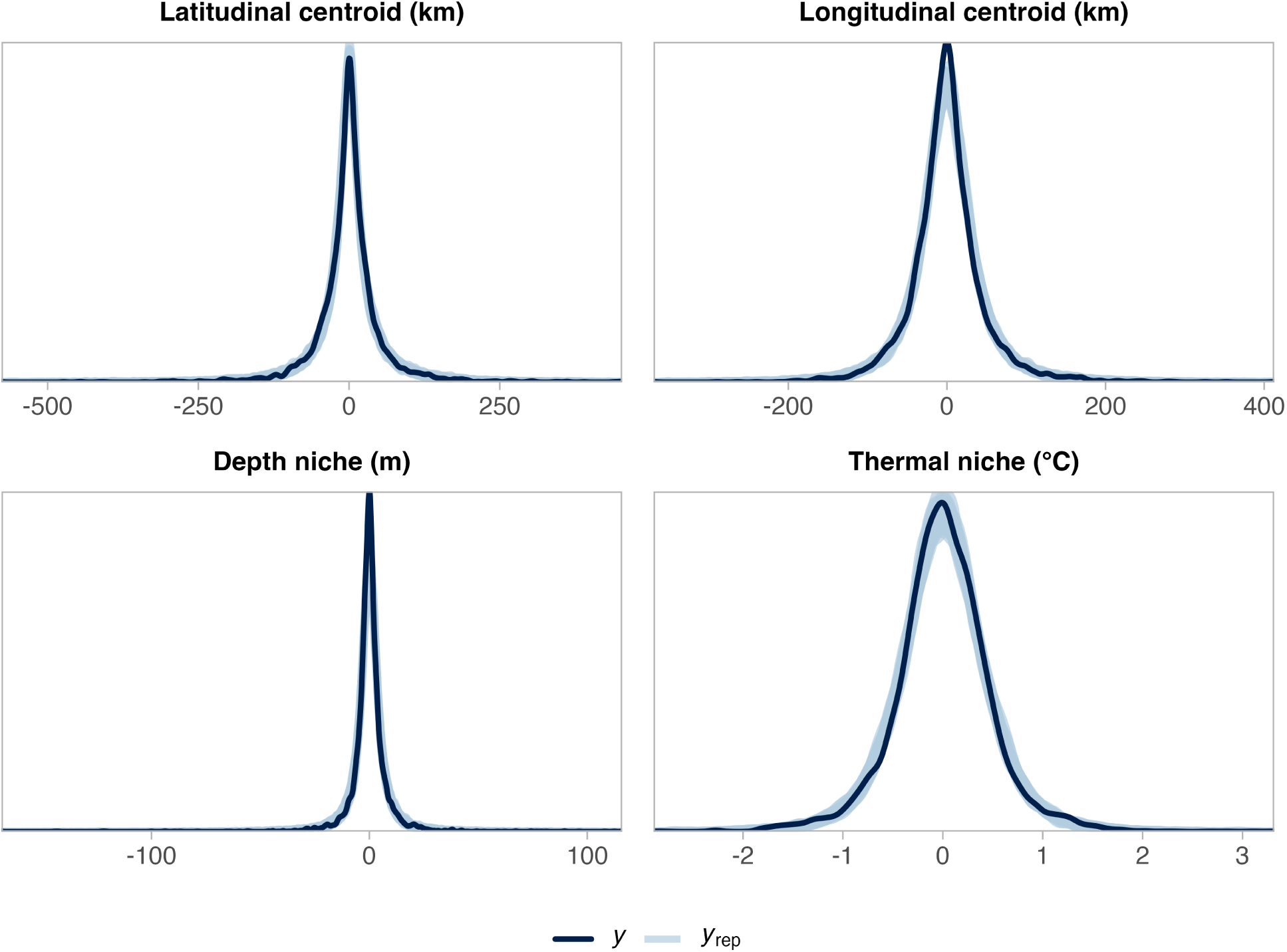
Posterior predictive checks for the four outcomes. Observed values (*y*) are compared with simulated datasets drawn from the posterior predictive distribution (*y_rep_*). Close alignment between the two indicates the model is capturing key features of the data generating process.

**Fig. B4:**
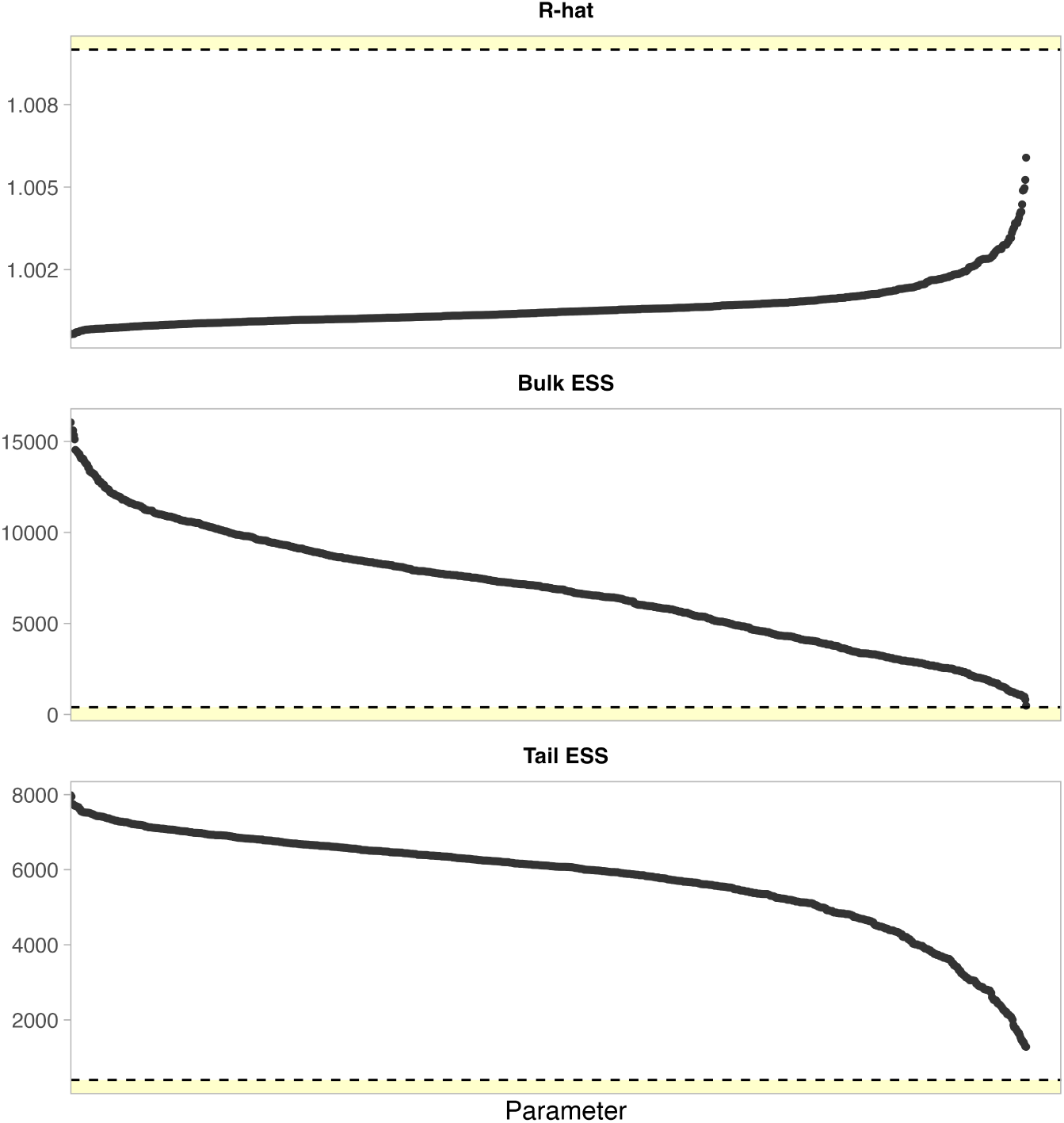
Convergence diagnostics for the hierarchical trend model. Panels show *R*^^^, bulk effective sample size (ESS), and tail ESS for all parameters. Yellow regions indicate values outside recommended thresholds.

